# Detection of NDM-1, VIM-1 and AIM-type metallo-beta-lactamase genes in Gram-negative bacteria isolated from clinical samples in Tamil Nadu

**DOI:** 10.1101/2020.11.29.403220

**Authors:** Prasanth Manohar, Aemy Joseph, B Karthika, Pradeep AnuPriya, Swetha S Mani, VS Varsha, Nachimuthu Ramesh

## Abstract

The distribution of carbapenem-resistant Gram-negative bacteria has become an increasing public health concern in India. The aim of this study was to investigate the prevalence of carbapenem-resistant bacteria isolated from the clinical samples in Tamil Nadu, India. A total of 126 non-repetitive Gram-negative bacteria were taken for this study. The susceptibility to meropenem was determined by Minimum Inhibitory Concentration (MIC) by broth micro-dilution. The phenotypic resistance screening such as MHT (Modified Hodge test), EDTA disk synergy and CIM (carbapenem inactivation method) were performed. A multiplex PCR was used for the detection of carbapenemase-encoding genes. Among the 126 isolates studied, 82 (65.07%) meropenem-resistant isolates were identified by MIC. A total of 18 (21.9%) isolates were found to be positive for Metallo-β-Lactamase production through EDTA synergy test. None of the isolates were carbapenemase producer by MHT and CIM. The isolates identified with resistance genes (8/82) were *bla*_NDM-1_ in two *Klebsiella* sp., two *P. aeruginosa* and one *A. baumannii*, *bla*_VIM-1_ in one *P. aeruginosa* and *bla*_AIM-1_ in one *P. aeruginosa* and one *A. baumannii*. The study showed the distribution and increase of carbapenem-resistant bacteria in the study region. Therefore, constant monitoring and effective elimination should be focused to reduce the spread of carbapenem-resistant isolates.

## Introduction

Antibiotic resistance has become a global threat with the rampant use of antibiotics especially in the developing countries like India. One of the important reasons for resistance development is the misuse of antibiotics, especially most of the clinicians prefer treatment based on patient symptoms, whereas very few microbiological samples are taken for culture and antibiotic sensitivity testing^1^. Gram-negative bacterial infections are wide spread, particularly, infections caused by multi-drug resistant *Enterobacteriaceae*, *Pseudomonas* and *Acinetobacter* have left us with the minimal therapeutic options. The Gram-negative bacterial infections are treated using broad-spectrum antibiotics and carbapenem is used as the last-resort for the hospital-acquired infections, especially, in intensive care units (ICU)^2,3^

Recently, resistance to carbapenem such as ertapenem, meropenem and imipenem was exhibited by *Enterobacteriaceae* called as carbapenem-resistant *Enterobacteriaceae* (CRE). Resistance to carbapenem can be brought by various mechanisms including the β-lactam hydrolysing enzyme such as carbapenemase, over expression of efflux pump, lack of porins in the bacterial cell membrane and the inability of a drug to bind with penicillin-binding protein^4^. The multi-drug resistant *Enterobacteriaceae* and non-fermenting Gram-negative bacteria are capable of genetically transmitting the resistance genes among the bacterial populations through transposons, plasmids and integrons leading to the exponential spread of such bacteria^4,5^. Genes encoding carbapenemase enzyme expand swiftly with different bacterial species through horizontal gene transfer (HGT). The resistance genes include KPC, GES/IBC, SME, NMC-A, IMI and SFC of Ambler class A and NDM, VIM, IMP, SPM, GIM, SIM, KHM,AIM, DIM, SMB, TMB and FIM Metallo-beta-lactamases of Ambler class B^6,7^. Since these genes are located on mobile genetic elements, it can spread widely, particularly, the genes NDM-1- and OXA-48-like which is very common in India^8,9,10,11^.

This suggests a serious threat which is imposed to public health due to the emergence and dissemination of carbapenem-resistant bacteria in the recent times and its association with high mortality and the potential to spread widely^12^. This avails for a rapid and accurate routine protocol for resistant bacterial screening and detection. This is vital in patient care and infection control in order to institute correct, targeted treatment and to reduce the escalation of resistance^13^. The U.S Centres for Disease Control and Prevention (CDC) has declared carbapenem-resistant bacteria as a major clinical threat. Therefore, control measures have to be taken to trace the dissemination of resistance genes. The primary objective of this study was to evaluate the distribution of carbapenem-resistant bacterial isolates from clinical samples collected from Tamil Nadu.

## Materials and methods

### Bacterial isolates

A total of 126 non-repetitive, Gram-negative isolates that were isolated from clinical samples (which included urine, pus, sputum, left ear swab. right ear swab, bronchial aspirate) from diagnostic centres in Chennai, Tamil Nadu were taken for this study. The clinical isolates were received at Antibiotic Resistance and Phage Therapy Laboratory, VIT and all the isolates were stored at −20°C until processing. The bacterial identification was carried out using VITEK identification system.

### Antibiotic Susceptibility Testing (AST)

Kirby Bauer Disc Diffusion method was performed to determine antibiotic susceptibility pattern. The bacterial suspension was adjusted to 0.5 McFarland turbidity standards and swabbed on to Muller Hinton Agar (MHA) plates (Hi-Media, India). The plates were briefly dried and antibiotic impregnated discs of meropenem (10 mcg) were placed on the surface of the agar and incubated at 37°C for 18 hours. The diameter of zones of inhibition was measured and interpreted using the CLSI guidelines^14^.

### Minimum Inhibitory Concentration (MIC)

MICs of meropenem were determined by standard micro-broth dilution method using 96 well micro-titre plates. The meropenem (10 mg/ml) was serially diluted in Muller Hinton No.2 cation adjusted broth (Hi-Media, India) to obtain a two-fold dilution series ranging from 0.125 μg/ml to 128 μg/ml^15^. A 5 μl of bacterial culture at 0.5 McFarland turbidity standards was added to the wells and plates were incubated for 18 hours at 37°C. The susceptibility profiles were determined by comparing the MIC values with breakpoints of each isolates recommended by CLSI guidelines^16^.

### EDTA-Disc Synergy Test

EDTA-Disc Synergy Test (Double Disc Synergy Test) was performed to determine the Metallo-β-lactamase (MBL) production by bacterial isolates. The MHA plates were swabbed with the bacterial isolates at 0.5 McFarland turbidity standards. After brief drying, two 10 μg meropenem disks were placed 20 mm apart, and 10 μl of 0.5 M EDTA solution (Hi-Media) was added to one of the disks. Zone enhancement in the area between two discs was evaluated as a positive result^17^.

### Modified Hodge Test (MHT)

Modified Hodge Test was performed for the detection of carbapenemase production. A lawn culture of the indicator organism, *E. coli* DH5α was made on MHA plates and allowed to dry for 3-5 minutes. A 10 μg meropenem disc was placed at the centre of the plate^18^. Carbapenem-resistant bacterial isolates from an overnight culture were streaked (single line) from the edge of the disc to the periphery of the plate. The plates were incubated at 37°C for 18 hours. Carbapenemase production was detected by a clover-leaf indentation of indicator organism towards meropenem disc^19^.

### Carbapenem Inactivation Method (CIM)

A suspension was made by homogenizing the culture in 1 ml of MH broth. Subsequently, 10 μg meropenem discs was added to the bacterial suspension and incubated for a minimum of 2 hours at 37°C. After the incubation, disc was placed on MHA plate inoculated with indicator strain *E. coli* DH5α (at McFarland standards value of 0.5) and further incubated at 37°C for 24 hours^20^. The absence of an inhibition zone was interpreted as the inactivation of meropenem in the susceptibility disc by carbapenemase produced by the bacterial isolates whereas; a clear inhibition zone indicated the absence of carbapenemase activity.

### DNA Isolation

DNA isolation was performed by boiling preparation method. Overnight grown cultures were centrifuged at 8000 x g for 10 minutes. 100 μl of sterile distilled water was added to the harvested bacterial cells and the cells were boiled for 15 min. The mixture was centrifuged at 2000 x g for 2 min. The extracted bacterial DNA in the supernatant was transferred to a sterile tube and stored at −20°C until molecular analysis^21^.

### Screening of antibiotic resistance gene determinants

Multiplex PCR was performed to detect the presence of β-lactamase genes *bla*_KPC,_ *bla*_IMP,_ *bla*_VIM-1,_ *bla*_NDM-1,_ *bla*_OXA-48-like,_ *bla*_AIM,_ *bla*_GIM,_ *bla*_BIC,_ *bla*_SIM_ and *bla*_DIM_^22,23^. The PCR products were sequenced at Eurofins analytical Services India Pvt Ltd Bangalore. The nucleotide blast (NCBI) was employed to compare the sequences with the GenBank sequences (www.ncbi.nlm.nih.gov/BLAST)^24^. The sequences were submitted to NCBI using BankIt tool and accession numbers were obtained.

## Results

### Carbapenem-resistant isolates

For this study, *Pseudomonas aeruginosa* (n=37), *Klebsiella* sp. (n=33), *Escherichia coli* (n=31), *Enterobacter* sp. (n=13), *Proteus* sp. (n=10) and *Acinetobacter baumannii* (n=2) were taken. Of the 126 isolates, 65.07% (n=82) were found to be resistant to meropenem by MIC (Fig.1). Out of which 41.4% (n=34) belonged to *P. aeruginosa* and 69.6% (n=23) belonged to *K. pneumoniae*. Interestingly, 56/126 (44.4%) isolates were found to be resistant to meropenem by disc diffusion test, whereas, three (5.08%) isolates exhibited intermediate resistance.

**Figure 1:**
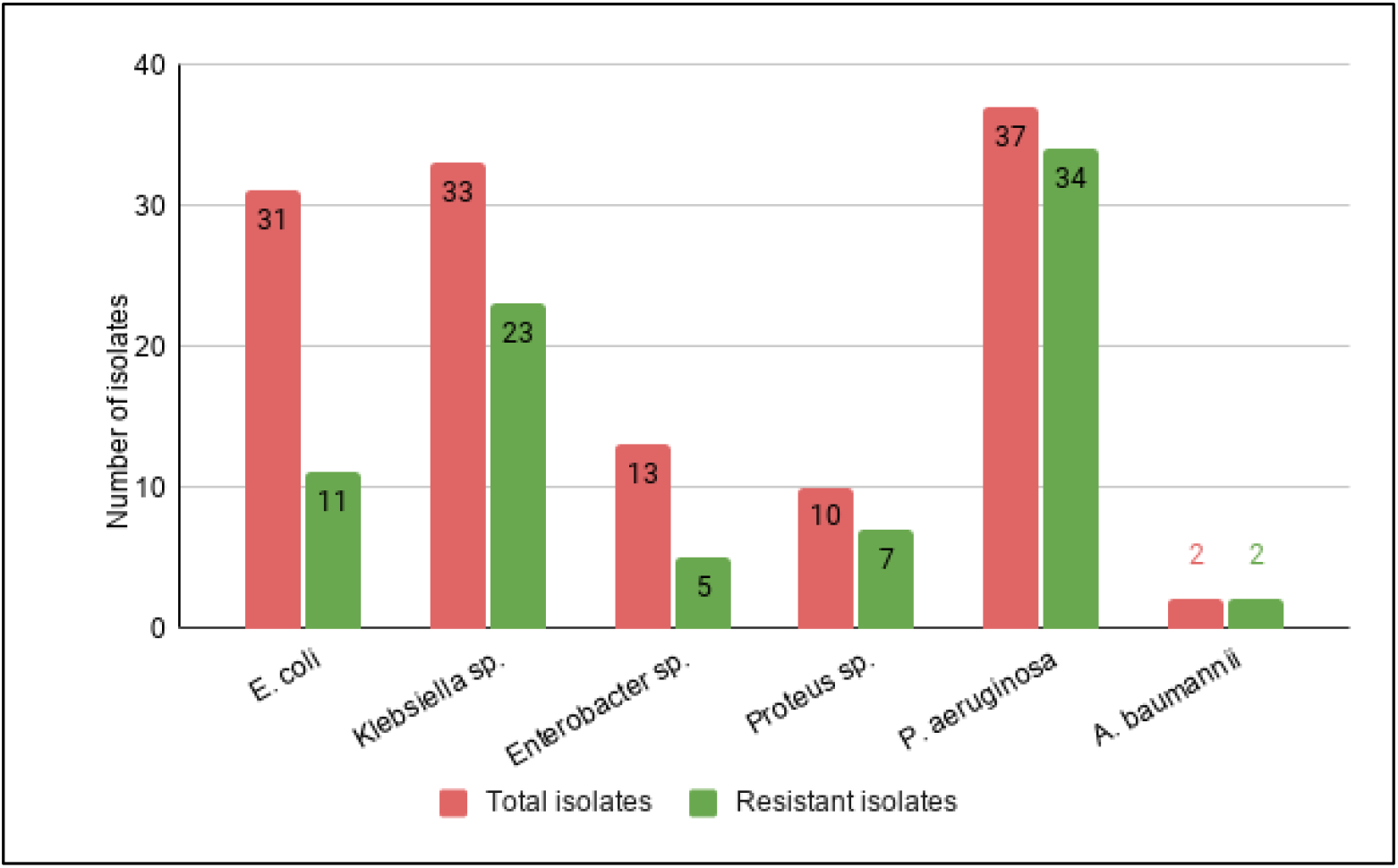
The number of meropenem-resistant isolates based on the MIC results.

### MBL or Carbapenemase producers

The MBL-producers and non-MBL-producers were distinguished using EDTA synergy test. Of the tested isolates (n=82), enhanced inhibition zones were exhibited by 18 (21.9%) isolates that were carbapenem-resistant. The presence of carbapenemase production in the isolates was determined by two methods, MHT and CIM. Among the 82 isolates, none of the isolates were found to be producing carbapenemase enzyme by MHT. All the 82 isolates showed negative results for CIM indicating the absence of carbapenemase production which further validated the results of MHT.

### Molecular studies

Total DNA was extracted from 82 resistant isolates and 10 carbapenem-resistance genes were screened. It was found that 8/82 isolates were harbouring at least one of the tested carbapenemase genes. The five isolates, 2 *Klebsiella* sp., 2 *P. aeruginosa* and 1 *A. baumannii* showed the presence of *bla*_NDM-1_ carbapenemase gene. One *P. aeruginosa* had *bla*_VIM-1_, and *bla*_AIM_-type was detected in one *P. aeruginosa* and one *A. baumannii* (Table 1). Following sequencing and BLAST analysis, the identified gene sequence showed 100% sequence similarities with varied strains of *bla*_NDM-1_ variant. The sequences were deposited under the accession number MK649943, MK649944 (for *Klebsiella*), and MK652488, MK652489 (for *P. aeruginosa*). Presence of multiple carbapenemase genes were not observed in the isolates.

**Table 1:** List of resistant isolates carrying carbapenem-resistance genes.

| Bacteria | *bla*<sub>NDM-1</sub> | *bla*<sub>VIM-1</sub> | *bla*<sub>AIM-type</sub> |
| --- | --- | --- | --- |
| *Klebsiella pneumoniae* | 1 | - | - |
| *Klebsiella oxytoca* | 1 | - | - |
| *Pseudomonas aeruginosa* | 2 | 1 | 1 |
| *Acinetobacter baumannii* | 1 | - | 1 |

## Discussion

From the study, the prevalence of carbapenem-resistant Gram-negative bacteria from the clinical sample was found to be 65% which emphasizes the need for controlling the dissemination of carbapenem-resistant bacteria. *P. aeruginosa* is one among the carbapenem-resistant bacteria tested in this study. Our previous studies also insisted the prevalence and dissemination of antibiotic resistant bacteria in Tamil Nadu, particularly the isolates carrying CTX-M^25^, NDM^8,9^, and OXA^8,9^. The CRE infected patients must be monitored and isolated as they may serve as a pool for the spread of infection and contaminating the environment^26^. Predominantly, the presence of carbapenem-resistance is estimated based on the MIC results where meropenem serves as an antibiotic of choice (in India) for screening of resistant isolates. It helps in the selection of antibiotic therapy and for guiding the appropriate dosing regimen.

Highly distinguishing data was observed for carbapenem-resistance between disc diffusion test and broth dilution method. In this study, 44.4% isolates exhibited resistance by disc diffusion, whereas, 65.07% by broth dilution method. The data suggests that the disc diffusion method is not valid for confirming the susceptibility; the values were not significant and also displayed contrasting results with lesser reproducibility. Thus, disc diffusion can be a qualitative measure; confirmatory results should be validated through MIC determination. The MIC tests can be used for detecting the resistant isolates and are reproducible.

As carbapenemase producing genes are transferred widely, a standard laboratory assay is required for tracing. MHT is a sensitive method for screening carbapenemase production but can infer false positive results with those strains having reduced porins or with extended spectrum beta-lactamases production^27^. Here, all the isolates showed negative results indicating some other mechanism or beta-lactamases is responsible for carbapenem-resistance. The strains with reduced MIC for carbapenem can be screened for carbapenemase by CIM. It also allows for the differentiation in resistance mechanisms i.e. through decreased permeability or due to beta-lactam hydrolysing enzymes. Thus, allowing the screening for extended-spectrum beta-lactamase production. Of the 82 resistant isolates, none of the isolates were positive by CIM. The metallo-β-lactamases production was screened for the isolates by performing EDTA synergy test. EDTA chelates Zn ions of the carbapenemase resulting in inactivation of the enzyme, thereby the diameter of inhibition zone increases^28^. The study showed that most of the resistant isolates determined through MIC showed the presence of metallo-β-lactamases. These strains are associated with high death rate and can hydrolyse lactamases and pose high threat during infection. The responsiveness and accuracy of gene detection can be enhanced by combined phenotypic detection by MHT and EDTA synergy^29^. Earlier studies performed by the authors in South India showed the predominant presence of carbapenem-resistance genes, NDM and OXA-48^8,9^. Here, the screening for resistance genes showed the presence of *bla*_NDM-1_ in *K. pneumoniae*, *P. aeruginosa* and *A. baumannii* which is Amber class B beta-lactamases responsible for the production carbapenemase enzyme. But it was not identified in our phenotypic studies. The presence of *bla*_VIM-1_ in *P. aeruginosa* becomes very common in India which clearly shows the dissemination of resistance within the Indian population. Both *P. aeruginosa* and *A. baumannii* carrying *bla*_AIM_-type carbapenemase is very uncommon in South India which needs further epidemiological reports. Proper monitoring and early detection of resistance genes is necessary as they can lead to increasing MDR infections. Also these genes may be associated with other resistome and cause rapid outbreak.

Since these molecular methods are slow and costly, laboratory diagnosis can make use of MIC detection for early diagnosis. Thus, awareness can be made among the public about hygiene practices and proper disposal of waste and controlled use of antibiotics. Since carbapenem-resistant bacteria are rapidly adapting and surviving through various mechanisms, alternative therapy is in high demand. With the increasing resistance reports, combination therapy has been chosen as an alternative. Accordingly, previous studies showed the synergistic activity between β-lactams and quinolones against Gram-negative and Gram-positive bacteria^30,31^. Based on the *in vitro* studies, ciprofloxacin appears to be a potent antibiotic when used in combination with meropenem against carbapenem-resistant isolates.

In conclusion, the carbapenemase enzyme distribution is imposing serious threat to antibiotic treatment. Any immediate measures have to be taken to prevent the emerging resistant strains and strategies for controlling the hospital-acquired infections and its implementation. The current instrument based molecular techniques and treatment should be improved for a rapid and effective screening and diagnosis particularly for the resistance genes in bacteria.

## Acknowledgement

The authors thank VIT for providing ‘VIT SEED GRANT’ for carrying out this research work.

## REFERENCES

1. Gould, I. M. International Journal of Antimicrobial Agents The epidemiology of antibiotic resistance. 2–9 (2008). doi:10.1016/j.ijantimicag.2008.06.016

2. Pawar, S. K., Patil, S. R., Karande, G. S., Mohite, S. T. & Pawar, V. S. Antimicrobial Sensitivity Pattern of Clinical Isolates in Intensive Care Unit in a Tertiary Care Hospital from Western India. 4, 108–113 (2016).

3. Manuscript, A. NIH Public Access. 80, 225–233 (2014).

4. Meletis, G. Carbapenem-resistance: overview of the problem and future perspectives. 15–21 (2016). doi:10.1177/2049936115621709

5. Agarwal, S., Kakati, B., Khanduri, S. & Gupta, S. Emergence of Carbapenem-resistant Non-fermenting Gram-Negative Bacilli Isolated in an Icu of a Tertiary Care Hospital. 11, 12–15 (2017).

6. Woodford, N., Wareham, D. W., Guerra, B. & Teale, C. Carbapenemase-producing Enterobacteriaceae and non-Enterobacteriaceae from animals and the environment: an emerging public health risk of our own making? 287–291 (2014). doi:10.1093/jac/dkt392

7. Bonomo, R. A. “Stormy waters ahead”: global emergence of carbapenemases. 4, 1–17 (2013).

8. Manohar P, Shanthini T, Ayyanar R, Bozdogan B, Wilson A, Tamhankar AJ, Nachimuthu R, Lopes BS. The distribution of carbapenem-and colistin-resistance in Gram-negative bacteria from the Tamil Nadu region in India. Journal of medical microbiology. 2017 Jul 1;66(7):874–83.

9. Nachimuthu R, Subramani R, Maray S, Gothandam KM, Sivamangala K, Manohar P, Bozdogan B. Characterization of carbapenem-resistant Gram-negative bacteria from Tamil Nadu. Journal of Chemotherapy. 2016 Sep 2;28(5):371–4.

10. Shanthini T, Manohar P, Samna S, Srividya R, Bozdogan B, Rameshpathy M, Ramesh N. Emergence of plasmid-borne blaoxa-181 gene in Ochrobactrum intermedium: first report from India. Access Microbiology. 2019 May 1;1(3):e000024.

11. Manohar P, Ragavi M, Augustine A, Hrishikesh MV, Ramesh N. Identification of blaGIM-1 in Acinetobacter variabilis isolated from the hospital environment in Tamil Nadu, India. bioRxiv. 2019 Jan 1:586164.

12. Diseases, Z. I. Facility Guidance for Control of Carbapenem-resistant Enterobacteriaceae (CRE) November 2015 Update - CRE Toolkit. (2015).

13. Datta, P., Gupta, V., Garg, S. & Chander, J. Phenotypic method for differentiation of carbapenemases in Enterobacteriaceae: Study from north India. 55, (2012).

14. Genen, H. V. E. B. Prevalence of extended-spectrum beta-lactamase genes in Acinetobacter baumannii strains isolated from nosocomial infections in Tehran, Iran. 14, 1–8 (2019).

15. Mello, M. M. De et al. Antimicrobial photodynamic therapy against clinical isolates of carbapenem-susceptible and carbapenem-resistant Acinetobacter baumannii. (2019).

16. Bush, K., Dudley, M. N. & Hecht, D. W. Methods for Dilution Antimicrobial Susceptibility Tests for Bacteria That Grow Aerobically; Approved Standard — Eighth Edition. 29, (2009).

17. Galani, I. et al. Evaluation of different laboratory tests for the detection of metallo- b -lactamase production in Enterobacteriaceae. 548–553 (2008). doi:10.1093/jac/dkm535

18. Takayama, Y., Adachi, Y., Nihonyanagi, S. & Okamoto, R. Modified Hodge test using Mueller – Hinton agar supplemented with cloxacillin improves screening for carbapenemase-producing clinical isolates of Enterobacteriaceae. 1, 774–777 (2019).

19. Carvalhaes, C. G., Pica, R. C., Nicoletti, A. G., Xavier, D. E. & Gales, A. C. Cloverleaf test (modified Hodge test) for detecting carbapenemase production in Klebsiella pneumoniae: be aware of false positive results. 249–251 (2010). doi:10.1093/jac/dkp431

20. Zwaluw, K. Van Der, Haan, A. De, Pluister, G. N. & Bootsma, H. J. The Carbapenem Inactivation Method (CIM), a Simple and Low-Cost Alternative for the Carba NP Test to Assess Phenotypic Carbapenemase Activity in Gram-Negative Rods. 1–13 (2015). doi:10.1371/journal.pone.0123690

21. Article, O. Heat Treatment of Bacteria: A Simple Method of DNA Extraction for Molecular Techniques. 117–122 (2009).

22. Poirel, L., Walsh, T. R., Cuvillier, V. & Nordmann, P. Multiplex PCR for detection of acquired carbapenemase genes. Diagn. Microbiol. Infect. Dis. 70, 119–123 (2011).

23. Diag-, R. Laboratory Detection of Enterobacteriaceae That Produce and MBL Etest were often difficult to interpret. We recommend using molecular tests for the optimal detection of carbapen-. 50, 3877–3880 (2012).

24. Bahramian, A. et al. First report of New Delhi metallo-β-lactamase-6 (NDM-6) among Klebsiella pneumoniae ST147 strains isolated from dialysis patients in Iran. Infect. Genet. Evol. 6, #pagerange# (2019).

25. Nachimuthu R, Kannan VR, Bozdogan B, Krishnakumar V, Pandiyan S K, Manohar P. CTX-M-type ESBL-mediated resistance to third-generation cephalosporins and conjugative transfer of resistance in Gram-negative bacteria isolated from hospitals in Tamil Nadu, India. Access Microbiology. 2020 Jun 11:acmi000142.

26. Nagaraj, S., Chandran, S. P., Shamanna, P. & Macaden, R. Carbapenem-resistance among Escherichia coli and Klebsiella pneumoniae in a tertiary care hospital in south India. 30, 93–96 (2012).

27. Pasteran, F., Mendez, T., Rapoport, M., Guerriero, L. & Corso, A. Controlling False-Positive Results Obtained with the Hodge and Masuda Assays for Detection of Class A Carbapenemase in Species of Enterobacteriaceae by Incorporating Boronic Acid. 48, 1323–1332 (2010).

28. Miriagou, V., Tzelepi, E., Kotsakis, S. D. & Daikos, G. L. Combined disc methods for the detection of KPC- or VIM-positive Klebsiella pneumoniae: improving reliability for the double carbapenemase producers. (2013).

29. Rastegar, A., Azimi, L., Rahbar, M. & Fallah, F. Phenotypic detection of Klebsiella pneumoniae carbapenemase among burns patients: First report from Iran. Burns 39, 173–175 (2012).

30. Kelly, L. M. & Jacobs, M. R. Comparison of Agar Dilution, Microdilution, E-Test, and Disk Diffusion Methods for Testing Activity of Cefditoren against Streptococcus pneumoniae. 37, 3296–3299 (1999).

31. Methodology, T., Credito, K., Lin, G. & Appelbaum, P. C. Activity of Daptomycin Alone and in Combination with Rifampin and Gentamicin against Staphylococcus aureus Assessed by. 51, 1504–1507 (2007).

